# A pathogenic role for histone H3 copper reductase activity in a yeast model of Friedreich’s Ataxia

**DOI:** 10.1101/2021.06.14.448268

**Authors:** Oscar A. Campos, Narsis Attar, Nathan V. Mallipeddi, Chen Cheng, Maria Vogelauer, Stefan Schmollinger, Sabeeha S. Merchant, Siavash K. Kurdistani

## Abstract

Disruptions to iron-sulfur (Fe-S) clusters, essential cofactors for a broad range of proteins, cause widespread cellular defects resulting in human disease. An underappreciated source of damage to Fe-S clusters are cuprous (Cu^1+^) ions. Since histone H3 enzymatically produces Cu^1+^ to support copper-dependent functions, we asked whether this activity could become detrimental to Fe-S clusters. Here, we report that histone H3-mediated Cu^1+^ toxicity is a major determinant of cellular Fe-S cluster quotient. Inadequate Fe-S cluster supply, either due to diminished assembly as occurs in Friedreich’s Ataxia or defective distribution, causes severe metabolic and growth defects in *S. cerevisiae*. Decreasing Cu^1+^ abundance, through attenuation of histone cupric reductase activity or depletion of total cellular copper, restored Fe-S cluster-dependent metabolism and growth. Our findings reveal a novel interplay between chromatin and mitochondria in Fe-S cluster homeostasis, and a potential pathogenic role for histone enzyme activity and Cu^1+^ in diseases with Fe-S cluster dysfunction.

**Teaser:** Reduction of Cu^1+^ production by histone H3 restores cellular deficiencies caused by insufficient supply of iron-sulfur clusters.

## Introduction

The ancient and ubiquitous iron-sulfur (Fe-S) clusters (*1*) function as prosthetic groups in many enzymes in diverse metabolic and informational processes (*2*). The importance of Fe-S clusters to life is evidenced by the pathological consequences associated with their loss of function. For example, Friedreich’s Ataxia (FRDA) is a genetic, progressive, neurodegenerative disease in humans caused by mutations in the *FXN* gene encoding frataxin (*3*), a mitochondrial protein important for the assembly of Fe-S clusters (*4*).

The proteins involved in the cellular assembly of Fe-S clusters, which begins in mitochondria, are evolutionarily conserved across eukaryotes (*2, 5-7*). In the budding yeast, *Saccharomyces cerevisiae*, the homolog of frataxin, Yfh1, also plays an important role in Fe-S cluster assembly in mitochondria (*8, 9*). Loss of Yfh1 results in diminished Fe-S cluster abundance, impairing activities of Fe-S cluster dependent proteins and pathways. These include aconitase activity (*10, 11*) and syntheses of certain amino acids such as lysine and glutamate (*12, 13*).

The net supply of Fe-S clusters depends on the balance between production in the mitochondria and degradation at various downstream stages and locations. The need to maintain this supply is compounded by the vulnerability of Fe-S clusters to damage, including by oxidative stress (*14-16*) and reduced copper (Cu^1+^) ions (*17*). In addition to potentially causing oxidative stress (*18, 19*), the propensity of Cu^1+^ ions to interact with cysteine sulfhydryl groups with high affinity (*20*) allows Cu^1+^ to directly compete with Fe-S clusters for protein binding sites (*21*). Thus, Cu^1+^ ions can impair Fe-S cluster assembly, delivery, or stability in various enzymes (*21, 22*) even in anaerobic conditions (*17*). This notion is bolstered by the finding that overexpression of Yah1 in yeast, a protein critical for the assembly of the Fe-S clusters in the mitochondria, resulted in resistance to highly elevated copper abundance (*23*).

These considerations suggested to us that production of Fe-S clusters might be counter-balanced by Cu^1+^-induced damage, placing Fe-S cluster levels and Cu^1+^ abundance in a reciprocal equilibrium. We therefore reasoned that when Fe-S cluster assembly is impaired, such as in FRDA in humans or following the loss of Yfh1 in yeast, even physiological Cu^1+^ levels could become toxic as cells cannot maintain sufficient Fe-S cluster production to counteract Cu^1+^-induced damage. In turn, depletion of Cu^1+^ might mitigate the cellular dysfunction resulting from loss of Fe-S cluster assembly, effectively establishing a new equilibrium albeit at lower levels of both Fe-S clusters and Cu^1+^ (fig. S1).

We recently discovered that the eukaryotic histone H3 is a cupric reductase (*24*), catalyzing electron transfer and reduction of cupric ions (Cu^2+^) to Cu^1+^. In yeast, mutation of the histidine 113 residue of histone H3 to asparagine (*H3H113N*), a key Cu^2+^-binding residue, resulted in diminished Cu^1+^ abundance and impaired function of copper-dependent processes, such as mitochondrial respiration and superoxide dismutase function. Considering the potential toxicity of Cu^1+^, however, our findings raised the possibility that histone H3-mediated Cu^1+^ production could result in detrimental effects to the pool of Fe-S clusters, especially when Fe-S cluster supply is inadequate. Indeed, in this study, we found that the various cellular defects resulting from Yfh1 depletion are rescued by diminishing histone cupric reductase activity. Fe-S cluster dependent function is also recovered by total cellular copper depletion, supporting the conclusion that Cu^1+^ production by histone H3 impairs Fe-S cluster function. Our findings suggest that when Fe-S cluster production is compromised, the threshold for copper toxicity decreases, making the physiological levels of Cu^1+^ a liability. Reduction of Cu^1+^ levels, including inhibition of the histone H3 copper reductase activity, might therefore represent a novel therapeutic strategy for diseases caused by the dysfunction of Fe-S clusters, such as FRDA.

## Results

### Histone *H3H113N* mutation prevents the growth defects caused by disruption of Fe-S cluster biogenesis or distribution

The *H3H113N* mutation in yeast decreases the Cu^1+^ pool and diminishes copper utilization for copper-dependent processes (*24*). We reasoned that this same mutation might provide resistance against Cu^1+^ toxicity. To test this, we introduced the *H113N* mutation in the two chromosomal copies of the histone H3 gene to generate the yeast strain *H3*^*H113N*^. Despite its somewhat slower growth, this strain grew better than WT in fermentative media containing toxic levels of copper (Fig. 1A). We considered whether resistance to elevated copper depended on the metallothionein, Cup1, which plays a prominent role in limiting toxicity by sequestering excess intracellular Cu^1+^ (*25, 26*). However, *H3*^*H113N*^ still displayed substantial resistance to copper toxicity relative to WT when Cup1 was inactivated (*F8stop*) (fig. S2A). These results are consistent with decreased intracellular Cu^1+^ abundance in *H3*^*H113N*^ resulting in less damage by copper.

**Fig. 1.**
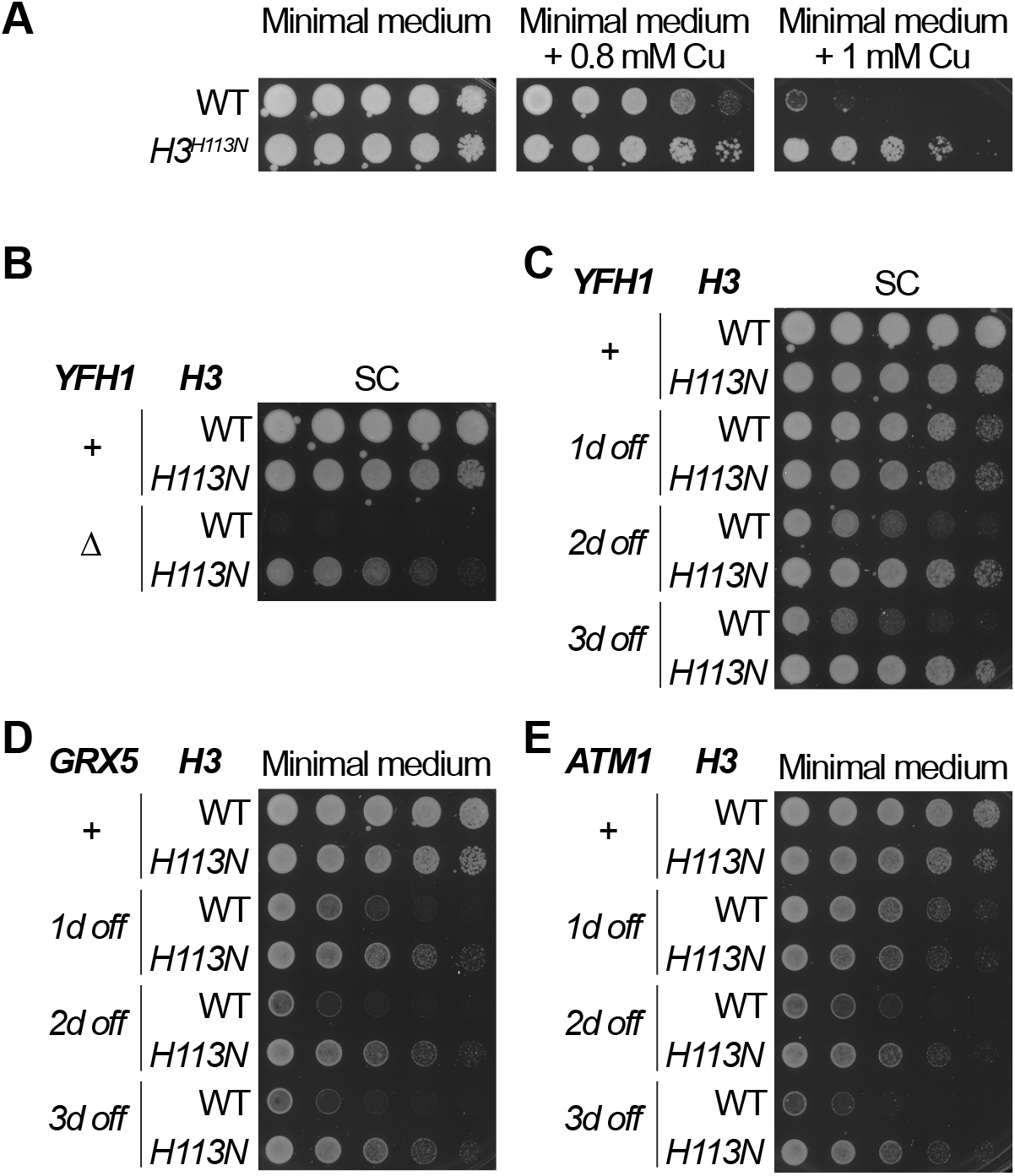
*H3H113N* rescues the growth defects caused by dysfunctional Fe-S cluster assembly or distribution. (**A**) Spot test assays in glucose-containing minimal media with or without additional CuSO_4_. Copper concentration in minimal medium at baseline is ∼0.25 µM. (**B**) Spot test assay in fermentative medium with or without *YFH1* deleted. (**C-E**) Spot tests with strains in which the *YFH1, GRX5*, or *ATM1* genes were placed under the control of the *GAL1* promoter, which is repressed in glucose-containing SC and minimal media. Genes were suppressed for 1, 2, or 3 days (d) prior to spot testing.

We next hypothesized that when Fe-S cluster assembly is compromised, diminishing histone cupric reductase activity, which decreases Cu^1+^ abundance (*24*), should mitigate the ensuing cellular defects, enabling cells to cope with decreased Fe-S cluster production (fig. S1). Deletion of most members of the mitochondrial Fe-S cluster assembly complex is lethal in yeast, except for *YFH1*. Deletion of *YFH1 (yfh1Δ*) results in severe growth defects in rich fermentative synthetic complete (SC) medium due to various underlying deficiencies (*8, 27*) (Fig. 1B). Importantly, the *H3H113N* mutation substantially rescued the severe growth defect of *yfh1Δ* (Fig. 1B).

Although *yfh1Δ* suffered severe growth defects, individual clones of this strain displayed a relatively high probability of spontaneous recovery of their growth defects over the course of a few days following *YFH1* deletion. This suggests a high likelihood of clones acquiring secondary mutations or other adaptations to cope with the disruption of Fe-S cluster homeostasis. To avoid this complication, we instead turned to a conditional transcriptional shutoff approach to deplete cells of Yfh1. We replaced the endogenous promoter of *YFH1* with the *GAL1* promoter, which is active in galactose-containing media but enables robust suppression of gene expression when cells are grown in the presence of glucose. This approach provides the additional benefit of enabling examination of the time-dependent deterioration of function when Yfh1 is depleted. Indeed, transcriptional shutoff of *YFH1* expression (*YFH1-off*) resulted in a minor growth defect after one day, but prolonged growth in the absence of *YFH1* expression revealed a gradually worsening defect in SC in two different yeast strains (Fig. 1C, fig. S2B, and Table S1). The growth defect of *YFH1-off* cells was exacerbated in media with substantially less amino acids (i.e., minimal medium) likely due to increased demand for Fe-S clusters for amino acid biosynthesis (fig. S2B and see below). Importantly, *H3H113N* substantially prevented this gradual growth deterioration in fermentative SC or minimal medium, consistent with diminished toxicity of Cu^1+^ to Fe-S clusters (Fig. 1C and fig. S2B).

To determine whether *H3H113N* could prevent the deterioration resulting from disruptions to Fe-S clusters more generally, we similarly repressed expression of either *GRX5* or *ATM1* with the *GAL1* promoter (*GRX5-off* and *ATM1-off*, respectively). Grx5 and Atm1 function downstream of Yfh1 in the transfer of Fe-S clusters to target proteins in the mitochondria and in the cytoplasm, respectively (*28-30*). Similar to the effect of Yfh1 depletion, *GRX5-off* and *ATM1-off* gradually deteriorated over time, but *H3H113N* substantially prevented this growth defect (Fig. 1D and 1E). These results suggest that *H3H113N* did not merely compensate for the specific loss of Yfh1, but instead, more generally prevented the dysfunction when Fe-S cluster homeostasis was disrupted.

### *H3H113N* prevents loss of Fe-S cluster-dependent amino acid synthesis

Depletion of Yfh1 may affect most, if not all, Fe-S cluster-dependent processes in the cell. These processes include the synthesis of certain amino acids that depend on enzymes containing Fe-S clusters (*12, 30-33*). All twenty amino acids are highly abundant in SC, which obviates the need for amino acid synthesis. We therefore gradually decreased the abundance of amino acids in SC and observed a corresponding gradual exacerbation of growth defects in the *YFH1-off* strain, even after a single day of Yfh1 shutoff (Fig. 2A and fig. S2B). This is consistent with an increased demand for Fe-S clusters for amino acid synthesis, thereby accentuating the deficiencies of *YFH1-off*. As in rich medium, the *H3H113N* mutation rescued the deterioration of growth in *YFH1-off* caused by amino acid limitation (Fig. 2A). With complete depletion of all amino acids, and thus when demand for Fe-S clusters would be highest, even *H3H113N* could not prevent the severe growth defect (Fig. 2A). This is likely because, even with decreased Cu^1+^-induced damage in *H3*^*H113N*^, total Fe-S supply remains below the threshold required to sustain sufficient amino acid synthesis.

**Fig. 2.**
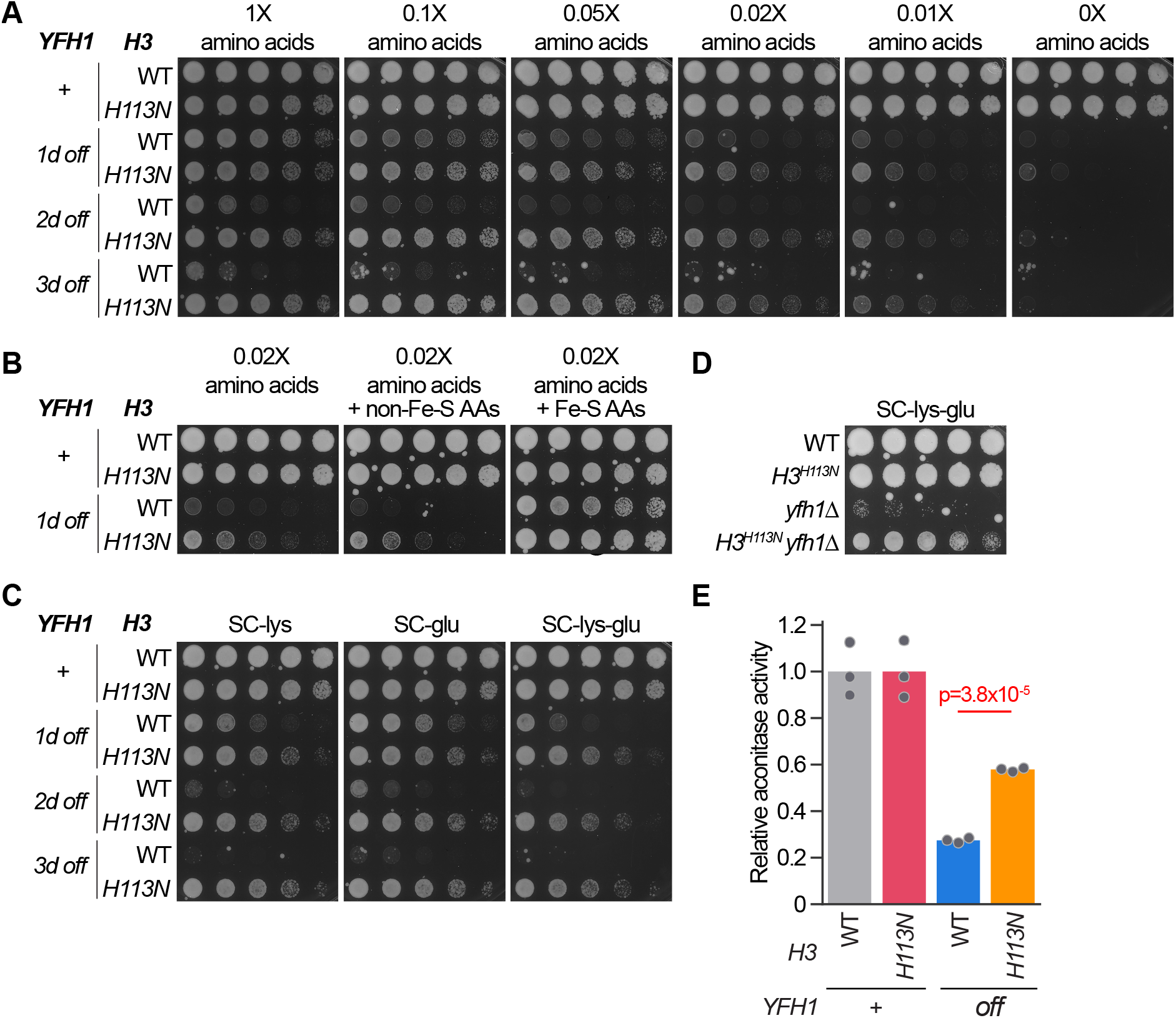
*H3H113N* restores Fe-S cluster-dependent amino acid synthesis. (**A**) Spot test assays in glucose-containing media with gradually decreasing amounts of all twenty amino acids. Baseline (i.e., 1X) concentration of each amino acid is 85.6 mg/L with some exceptions (see Methods). (**B**) Spot test assay with amino acid-depleted media ± 80 mg/L of each of the amino acids that are either not dependent (non-Fe-S AAs) or dependent (Fe-S AAs) on Fe-S cluster-containing enzymes. (**C and D)** Spot test assays with SC medium lacking lysine or glutamate. (**E**) Aconitase activity assay using whole cell extracts of the indicated strains grown in SC-lys-glu. *YFH1-off* strains were grown in SC medium for 44 hrs. Bars show mean activity relative to the matched WT or *H3*^*H113N*^, and each dot is an independent experiment (n = 3).

Fe-S cluster-containing enzymes participate in the syntheses of eight amino acids: methionine, cysteine, leucine, isoleucine, valine, lysine, glutamate, and glutamine. Syntheses of the others are not directly dependent on Fe-S clusters. Consistent with this, supplementation with eight amino acids, the syntheses of which are not dependent on Fe-S clusters, did not alleviate the growth defects of *YFH1-off*. In contrast, supplementation of amino acid-depleted media with only the eight Fe-S cluster-dependent amino acids restored growth of *YFH1-off* to the level displayed in complete medium (Fig. 2B). Altogether, these findings indicate that the growth defect of *YFH1-off* when amino acids were depleted was due to the diminished function of the Fe-S cluster-containing enzymes for synthesis of specific amino acids; and that diminishing copper reductase activity of histone H3 restores the function of these enzymes.

We further focused on two Fe-S cluster-dependent amino acids as testbeds for Fe-S cluster homeostasis. The removal of lysine and glutamate from otherwise replete medium (SC-lys-glu) substantially diminished growth of the *YFH1-off* strain almost as much as 100-fold depletion of all twenty amino acids (Fig. 2C). This confirms that lysine and glutamate syntheses alone placed increased demands for Fe-S clusters and exacerbated the growth defect of the *YFH1-off* strain (Fig. 2C). Similar to other conditions, decreased Cu^1+^ production in *H3*^*H113N*^ substantially prevented severe growth defects due to reduced lysine and glutamate syntheses resulting from diminishment (Fig. 2C) or complete loss of Yfh1 (Fig. 2D).

### *H3H113N* protects aconitase activity when Yfh1 is depleted

The ancient and conserved mitochondrial aconitase catalyzes the isomerization of citrate to isocitrate in the tricarboxylic acid, which in turn contributes to glutamate synthesis (*13*). Importantly, aconitase uses an Fe-S cluster cofactor for its catalytic function (*34-36*). Deficiencies in aconitase activity have been identified in heart biopsies of patients with FRDA, and similar deficiencies were identified in yeast strains lacking *YFH1* (*10*). We therefore assessed aconitase activity in whole cell extracts. Consistent with the growth defect of *YFH1-off* in SC-lys-glu, aconitase activity was significantly diminished following Yfh1 depletion (Fig. 2E). Furthermore, *H3H113N* significantly restored aconitase activity to ∼60% of normal levels, corroborating the rescue of the growth and amino acid synthesis defects (Fig. 2E).

### *H3H113N* prevents global transcriptional rewiring in response to defects in Fe-S cluster production

Yfh1 depletion leads to widespread rewiring of gene expression, including induction of an iron deficiency response (*37, 38*). We assessed gene expression profiles by mRNA sequencing and found substantial and reproducible alterations in *YFH1-off* in rich fermentative medium (Fig. 3A). Approximately half of the yeast genome (2867 genes) was significantly differentially expressed in *YFH1-off* compared to WT (Fig. 3B). However, *H3H113N* largely prevented the majority of gene expression changes, consistent with the ability to rescue the growth and amino acid synthesis defects resulting from Yfh1 and Fe-S cluster depletion (Fig. 3B). For example, genes that were at least 2-fold significantly upregulated (n=280) (Fig. 3C) or downregulated (n=214) (Fig. 3D) in the *YFH1-off* strain were largely restored to their normal expression levels in *H3*^*H113N*^ *YFH1-off*.

**Fig. 3.**
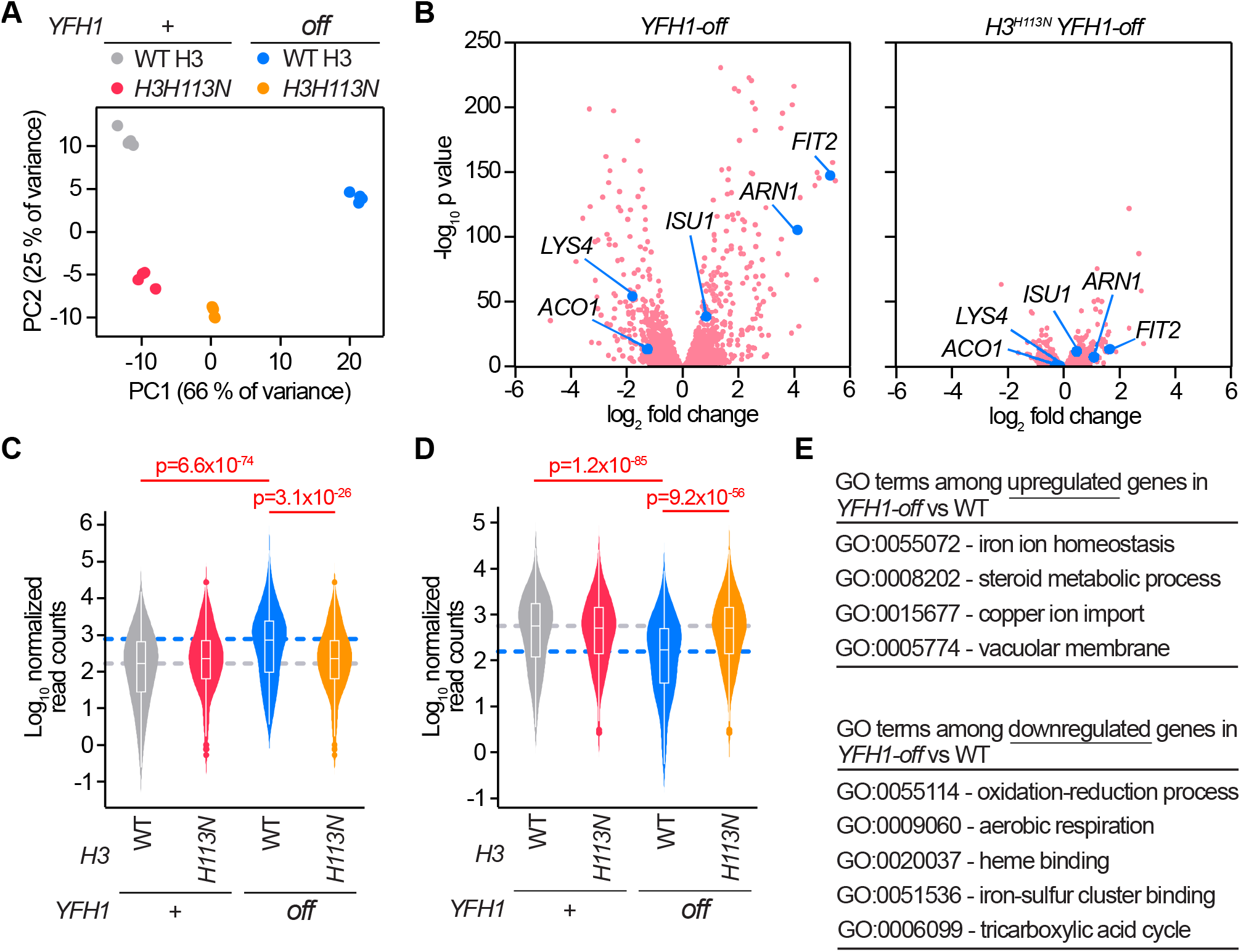
*H3H113N* prevents global transcriptional rewiring in *YFH1-off*. (**A**) Principal component (PC) analysis of global gene expression values from cells in SC medium from four independent experiments. *YFH1-off* strains were grown in SC medium for 44 hrs prior to RNA sequencing. (**B**) Volcano plots of average log2 fold changes of *YFH1-off* compared to WT (left) or *H3*^*H113N*^ *YFH1-off* compared to *H3*^*H113N*^ (right). A subset of iron regulon genes (*ARN1, FIT2*) or Fe-S cluster-binding genes (*ACO1, ISU1, LYS4*) are indicated. (**C and D**) Average mRNA expression levels for either (C) genes upregulated by at least 2-fold (n=280) or (D) genes downregulation by at least 2-fold (n=214) in *YFH1-off* compared to WT. The white box and whisker plots overlaid on the violin plots are the median and interquartile ranges. Dots are outlier data points. The grey and blue dashed lines are the median expression levels in WT and *YFH1-off*, respectively. (**E**) Significantly enriched gene ontology (GO) terms among the significantly differentially expressed genes in *YFH1-off* compared to WT.

As expected, there was significant enrichment for genes involved in iron homeostasis, uptake, and transport among the significantly upregulated genes in *YFH1-off* compared to WT (*37*) (Fig. 3E). This is consistent with the function of the transcription factor Aft1, which activates many genes in response to the loss of Fe-S clusters (*39, 40*). Correspondingly, among significantly downregulated genes in *YFH1-off* compared to WT, cellular processes that utilize Fe-S clusters as cofactors, including the mitochondrial respiratory chain, tricarboxylic acid cycle, and amino acid synthesis were significantly enriched (Fig. 3E). This finding is also consistent with previous investigations demonstrating that mRNA abundances of Fe-S cluster genes is generally diminished in concurrence with decreases in cellular iron abundance (*41, 42*). Altogether, the fact that *H3H113N* prevented most of the transcriptional changes in *YFH1-off*, as opposed to specific subsets of genes, such as the lysine and glutamate synthesis genes, is consistent with *H3H113N* broadly preventing disruptions to Fe-S clusters instead of rescuing specific downstream processes.

We considered more closely the expression of genes involved in iron mobilization and utilization, as well as the various Fe-S cluster assembly and target proteins. A majority of the Aft1 target genes displayed significantly elevated expression when Yfh1 was depleted (Fig. 3B and fig. S3A), which is consistent with the enrichment of this set of genes among all upregulated genes in *YFH1-off* compared to WT (Fig. 3E). The most highly induced Aft1 target genes were the cell wall mannoproteins *FIT1, FIT2*, and *FIT3*, as well as *ARN1* and *ARN2*, which together mediate uptake of siderophore-iron chelates from the environment (*43, 44*). Importantly, *H3H113N* ameliorated almost all of the changes resulting from Yfh1 depletion (Fig. 3B and fig. S3A). As Aft1 responds to the abundance of Fe-S clusters (*39*), this finding further suggests that *H3H113N* prevented Fe-S cluster disruptions.

Many of the genes encoding for Fe-S cluster binding proteins were also differentially expressed in response to Yfh1 depletion (Fig. 3B and fig. S3B). Several members of the mitochondrial Fe-S cluster assembly complex were upregulated, including *ISU1, ISU2*, and *NFS1* (Fig. 3B and fig. S3B). Induction of these genes suggests a response by cells to compensate for diminished Fe-S cluster assembly in mitochondria. Importantly, *H3H113N* almost entirely prevented the upregulation of the Fe-S cluster assembly proteins in response to Yfh1 depletion (Fig. 3B and fig. S3B), indicating that the histone H3 mutation does not rescue *YFH1-off* by enhancing the process of Fe-S cluster assembly. Furthermore, genes encoding most other Fe-S cluster binding proteins displayed significant downregulation in response to Yfh1 depletion (Fig. 3B and fig. S3B), in correspondence with decreased cellular Fe-S cluster abundance (*41*). Importantly, *H3H113N* mostly prevented the downregulation of these genes (Fig. 3B and fig. S3B), further suggesting an ability to maintain functional Fe-S clusters despite a diminishment of Fe-S cluster assembly.

### Iron abundance and superoxide dismutase do not account for the protective effect of *H3H113N* when Fe-S cluster assembly is disrupted

Previous studies have identified excessive iron accumulation as a major damaging component when frataxin/Yfh1 function is compromised (*45-47*). Reduced abundance of cellular Fe-S clusters triggers the activation of Aft1, which in turn, increases iron uptake from the environment. However, due to ineffective Fe-S cluster assembly in the absence of Yfh1, iron uptake does not effectively restore Fe-S cluster abundance. Instead, iron accumulates in the cell, including in mitochondria, which is proposed to become toxic (*40*). To determine if *H3H113N* was restoring function in *YFH1-off* by preventing iron accumulation, we first generated additional strains in which *FET3* was deleted (*fet3Δ*). The Fet3 multicopper oxidase is part of the high affinity iron uptake complex and, as an Aft1 target gene, substantially contributes to iron uptake (*48-50*). Consistent with this, *fet3Δ* prevented the deterioration of growth of *YFH1-off* in both SC and SC-lys-glu media (Fig. 4A). However, in contrast to the effect of *H3H113N, fet3Δ* could not sustain growth of *YFH1-off* after several days on solid agar media (Fig. 4A) and did not rescue *YFH1-off* in liquid culture (Fig. 4B). More important, the *H3*^*H113N*^ *YFH1-off fet3Δ* triple mutant strain grew substantially more than *YFH1-off fet3Δ* in SC-lys-glu (Fig. 4A), which suggests that the *H3H113N* prevented the loss of function through a mechanism other than preventing iron uptake.

**Fig. 4.**
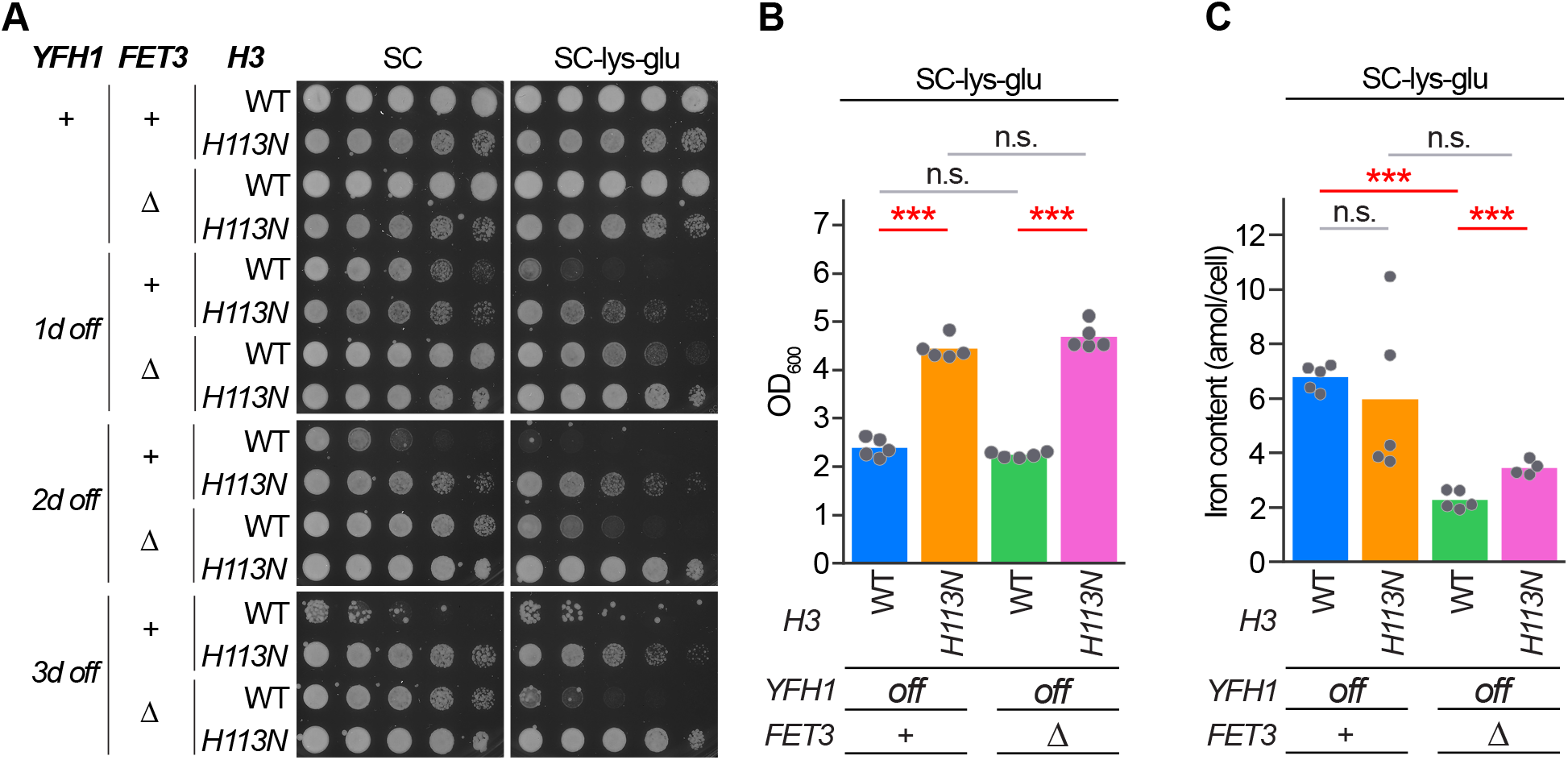
Differences in cellular iron content do not account for the protective effect of *H3H113N* when *YFH1* is shutoff. **(A)** Spot test assays in SC medium or SC without lysine and glutamate. **(B)** Growth after 44 hrs (see Methods for growth procedure) in liquid SC without lysine and glutamate. Bars show means and each dot is an independent experiment (n = 5). (**C**) Intracellular iron content for strains grown in liquid SC without lysine and glutamate. Bars show means and each dot is an independent experiment (n = 4-5). ***, p < 0.001; n.s., not significant.

To corroborate this finding, we examined cellular iron content in *YFH1-off* strains using inductively coupled plasma mass spectrometry. Yeast strains were grown in liquid media for the shortest amount of time required for Yfh1 depletion to result in a growth defect in SC-lys-glu medium (to avoid excessive cell death in the *YFH1-off* strain). This occurred after 44 hours of *YFH1* shutoff (Fig. 4B). Despite the rescue of *YFH1-off* growth defect by *H3H113N*, the cellular iron content in *YFH1-off* and *H3*^*H113N*^ *YFH1-off* were not significantly different (Fig. 4C). Furthermore, deletion of *FET3* decreased cellular iron content similarly in both strains (Fig. 4C). These findings therefore indicate that *H3H113N* rescues *YFH1-off* independently of changes in intracellular iron content.

Fe-S clusters are also highly susceptible to oxidative damage (*14, 16*). The Cu-Zn superoxide dismutase, Sod1, is a major contributor to antioxidant defense and is likely to protect Fe-S clusters from damage in the presence of oxygen (*51*). We previously found that *H3H113N* diminishes Sod1 activity (*24*). However, in the unique context of *YFH1-off* and the resulting decrease of Fe-S clusters, we considered whether the restoration of function by *H3H113N* depended on Sod1. We, therefore, generated additional strains in which *SOD1* was deleted (*sod1Δ*). Consistent with a protective function, *sod1Δ* exacerbated the deterioration of growth of *YFH1-off* in both SC and SC-glu media (fig. S4). Note that we did not use SC-lys-glu medium because *sod1Δ* is a lysine auxotroph (*52*). Critically, *H3H113N* was still able to restore function in the *YFH1-off sod1Δ* strain (fig. S4), indicating that the *H3H113N* protective effects are independent of superoxide dismutase activity.

### *H3H113N* diminishes Cu^1+^ toxicity to rescue Yfh1 depletion

Our finding that the *H3H113N* mutation results in resistance to copper toxicity (Fig. 1A), and that Cu^1+^ ions are especially toxic to Fe-S clusters, suggested that the broad rescue of *YFH1-off* by *H3H113N* was due to preventing copper toxicity. By the 44-hour time point, copper content had increased by ∼35% in *YFH1-off* compared to WT (Fig. 5A). This modest accumulation coincided with increased expression of copper mobilization and uptake genes, such as the main copper importer, *CTR1* (*53*), which supports iron homeostasis (*53*) and may be a direct target of Aft1 (*54*) (fig. S5A). Conversely, total copper abundance did not change in *H3*^*H113N*^ *YFH1-off* compared to *H3*^*H113N*^ (Fig. 5A). Addition of 10 µM copper in SC-lys-glu resulted in substantially increased total cellular copper content in *YFH1-off*, and even more so in *H3*^*H113N*^ *YFH1-off* (fig. S5B). Despite elevated copper accumulation, *H3H113N* still rescued the growth of *YFH1-off* in this media (fig. S5C). This indicates that the mechanism by which the *H3H113N* mutation restores Fe-S cluster homeostasis is not by diminishing total copper accumulation but rather by a specific decrease in Cu^1+^ abundance.

**Fig. 5.**
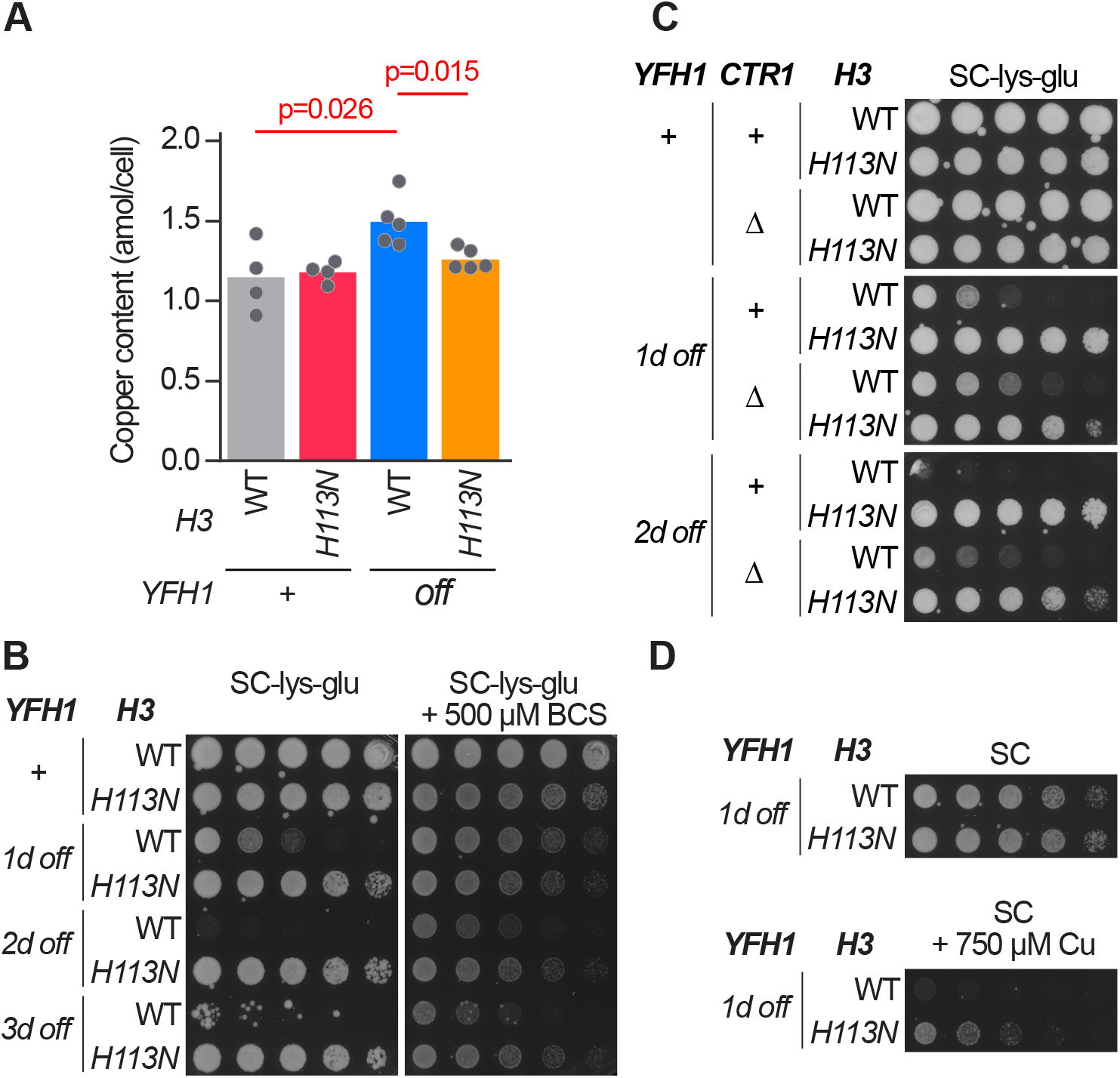
*H3H113N* rescues *YFH1-off* by diminishing copper toxicity. **(A)** Intracellular copper content for strains grown in liquid SC medium. *YFH1-off* strains were grown in SC for 44 hrs. Bars show means and each dot is an independent experiment (n = 4-5). (**B-D**) Spot test assays in SC medium or SC without lysine and glutamate and with or without the indicated amount of (B) BCS or (D) additional CuSO_4_.

To support the notion that *H3H113N* prevented Cu^1+^ toxicity when Yfh1 was diminished, we tested whether depletion of total cellular copper would perform similarly. Indeed, presence of the copper chelator bathocuproine disulfonate (BCS) rescued the growth of *YFH1-off* and *yfh1Δ* in SC-lys-glu medium, similar to but to a lesser extent than the effect of *H3H113N* (Fig. 5B and fig. S5D). Deletion of *CTR1* also prevented the defect in *YFH1-off* albeit less robustly after several days of growth of *YFH1-off* cells (fig. S5E). Importantly, the restoration of function by *H3H113N* was independent of *CTR1*, as the *H3*^*H113N*^ *YFH1-off ctr1Δ* triple mutant grew better than *YFH1-off ctr1Δ* (Fig. 5C). These findings suggest that the decrease in Cu^1+^ abundance specifically by *H3H113N* (*24*) is a distinct and more substantial protective effect for Fe-S clusters than merely diminishing total copper content. Finally, *H3H113N* ameliorated the increased sensitivity of *YFH1-off* to excessive exogenous copper (Fig. 5D), consistent with decreased Cu^1+^ abundance. Altogether, our findings demonstrate the robust ability of the *H3H113N* mutation to prevent the deterioration of cellular function when Fe-S cluster assembly is compromised.

## Discussion

Iron-sulfur clusters are essential enzymatic or structural cofactors for proteins in many important cellular processes in eukaryotes. The assembly of Fe-S clusters is initiated in mitochondria followed by distribution throughout the mitochondria, cytoplasm, and nucleus. These cofactors, however, are highly vulnerable to damage by Cu^1+^ ions through oxidative stress or direct displacement (*14, 17, 22*). Disruption of Fe-S cluster homeostasis has been implicated in the pathogenesis of several human diseases ranging from cancer to neurodegeneration (*3, 55-57*). Understanding the biological principles governing the production and decay of Fe-S clusters may therefore provide insights into human pathology and point to new therapeutic approaches.

In this study, we found that Cu^1+^ ions generated by the histone H3 cupric reductase activity function as key attrition factors against Fe-S clusters. This attrition could become toxic and result in metabolic and growth defects when Fe-S cluster assembly or distribution are compromised. Indeed, diminishment of Cu^1+^ by impairment of the cupric reductase capability of histone H3 (*24*) restored Fe-S cluster dependent functions when Fe-S cluster synthesis was inadequate—either due to decreased assembly (Figs. 1B and 1C) or defective transport (Figs. 1D and 1E). Our findings thus reveal that Fe-S cluster deficiencies due to a variety of insults can be restored by diminishing histone-mediated Cu^1+^ production. This presents Cu^1+^ depletion, for example, through inhibition of histone enzyme activity, as a new potential therapeutic modality when Fe-S cluster homeostasis is disrupted. Altogether, our work reveals a novel and opposing relationship between histone H3 and mitochondria in maintaining appropriate levels of Fe-S clusters in the cell. Alterations of this balance may contribute to disease pathology but could potentially be restored at a lower but functioning equilibrium to mitigate disease phenotypes (fig. S1).

While it is known that copper ions support iron homeostasis in eukaryotes, such as through enzymes like Fet3 in yeast or ceruloplasmin in humans, Cu^1+^ damages Fe-S clusters through oxidative stress and/or displacement. The precise mechanism by which the histone-dependent Cu^1+^ production in the nucleus can become toxic to the cell, including mitochondria, remains unclear. One possibility is that nuclear Cu^1+^ produced by histones that is normally trafficked to the mitochondria, in the context of disrupted Fe-S cluster assembly, damages the remaining co-factors in the mitochondria, in turn impairing downstream Fe-S dependent processes including amino acid synthesis. Another possibility is that histone-produced Cu^1+^ causes damage locally to Fe-S cluster proteins in the nucleus, such as the DNA polymerases and Rad3 (*58, 59*). To compensate, Fe-S clusters assembled elsewhere are redirected to replenish the various nuclear proteins but at the cost of compromising other Fe-S dependent processes. A third possibility is that Cu^1+^ toxicity is not limited to a subcellular location but occurs throughout the cell, with the loss of the most vulnerable Fe-S proteins manifesting as specific cellular phenotypes. Regardless of the mechanism of the damage, decreased histone H3 cupric reductase activity would diminish production of Cu^1+^, decreasing the possibility of damage to Fe-S clusters locally in the nucleus or at distant sites in the cytoplasm or mitochondria. Future investigations that trace the pathways of copper flux, distribution, and homeostasis should help reveal the mechanisms underlying the dynamics of Cu^1+^-induced damage to Fe-S clusters.

The fact that both histones and the mitochondrial Fe-S cluster assembly complex are highly conserved across eukaryotes suggests that the histone H3-mediated Cu^1+^ production may also become toxic in human disease with Fe-S cluster dysfunction. This prediction is especially relevant in diseases like FRDA and subtypes of sideroblastic anemia (*60, 61*). FRDA is a neurodegenerative disease caused by triplet expansion mutations in the first intron of frataxin, the human homolog of Yfh1 (*3*). These genetic perturbations diminish frataxin expression, resulting in increased mitochondrial iron, decreased activity of Fe-S cluster enzymes, and greater oxidative stress (*62*). Indeed, treatment with hypoxia can correct the various cellular defects resulting from frataxin loss and has been suggested as a potential therapeutic modality for FRDA (*63*). Sideroblastic anemia with spinocerebellar ataxia (ASAT) is caused by mutations in ABCB7, a mitochondrial ATP-binding cassette transporter and the human homolog of yeast Atm1, which can export Fe-S cluster intermediates from mitochondria (*64, 65*). At the cellular level, ASAT is characterized by aberrant mitochondrial iron homeostasis (*66*). Our findings that *H3H113N* could protect against the dysfunction resulting from depletion of either Yfh1 or Atm1 implicate histone-dependent Cu^1+^ toxicity as a possible pathogenic factor in FRDA and ASAT, and potentially other diseases based on disrupted iron homeostasis. Our findings in yeast, if confirmed in studies of mammalian systems, also suggest possible new therapeutic opportunities.

## Materials and Methods

### Strains and general growth conditions

*Saccharomyces cerevisiae* strains used in this study are based on the YLK1879 (S288C background, MATa) and BY4741 (S288C background, MATa) (*67*) strains, the former of which was a generous gift from Leonid Kruglyak (*68*). Strains are listed in Table S1. Strains were generally maintained on YPD (1% Yeast extract, 2% Peptone, 2% D-glucose) plates, which were supplemented with 20 µM CuSO4 for strains lacking *CTR1*. Strains with the *GAL1* promoter placed upstream of *YFH1, GRX5*, or *ATM1* were maintained on YPG (1% Yeast extract, 2% Peptone, 2% Galactose) plates. All strains were grown at 30°C in all experiments.

### Strain generation

The *H3H113N* mutation was generated in both chromosomal loci (*HHT1* and *HHT2*) using the CRISPR-Cas9 system optimized for *S. cerevisiae (69*). Subsequent gene deletions and promoter insertions were generated by standard yeast gene replacement and targeted insertion methodology using selectable marker integration (*70*). Successful integrations and deletions were confirmed by PCR. The *CUP1* gene was disrupted by introduction of a stop codon at Phe8 at all *CUP1* copies using the CRISPR-Cas9 system. Strains in which *YFH1* was deleted were phenotypically unstable. Within a few days following the transformation to replace *YFH1* with a selectable marker, clones would often spontaneously recover growth rates. Therefore, for the experiments shown in Figs. 1B, 2D, and figs. S5D, *yfh1Δ* strains were used immediately following transformation and not stored long-term. In all other cases, standard yeast culturing procedures were followed, and strains were continuously maintained on YPD or YPG plates, restreaking cells every two weeks for a maximum of 4 times.

### Preparation of solutions, media, and glassware

Removal of contaminating metals from glassware and solutions is a critical precaution to ensure reproducibility of experiments. All glassware was treated with 3.7% hydrochloric acid for ≥ 12 hours, followed in some cases by ≥ 12 hours with 7% nitric acid to remove trace metal contamination. All solutions, buffers, and washes were prepared using Milli-Q (Millipore Sigma) ultra-pure water. Solutions were prepared using BioUltra grade (Sigma) reagents, when available. For yeast media, addition of all components was done without the use of metal spatulas. Media were filtered through 0.2 µm membranes. Fermentative media were SC (synthetic complete medium with 2% glucose, all amino acids, and uracil and adenine), SC lacking lysine or glutamate, or minimal medium (with 2% glucose and either no amino acids, or only essential auxotrophic metabolites). Agar media were prepared using acid-washed glassware.

### Spot tests

Cells from exponentially growing cultures were 4-fold serially diluted and spotted on various media conditions as indicated in the figures. For experiments in which the *GAL1* promoter was used to shutoff expression for 1, 2, or 3 days, cells on YPG were inoculated into SC medium and grown for approximately 15 hours (day 1). For 2 and 3 days of shutoff, cultures were grown for an additional 48 hours in SC and diluted to OD_600_ = 0.05 every 24 hours. Cells were allowed to grow for an additional 4 hours to enter exponential growth phase, at which point they were collected for spotting. Spotted plates were incubated at 30°C for up to 4 days and imaged daily. Images shown in the figures were captured when sufficient growth had occurred and growth differences could be assessed (2-4 days). For the experiment in Fig. 2A, plates with gradually decreasing amounts of SC amino acid powder (Sunrise Science Products) were prepared, while keeping all other components the same (2% glucose, 1.7 g/L yeast nitrogen base, 5 g/L ammonium sulfate).

### Growth of *YFH1-off* strains in liquid culture

For aconitase assay, inductively coupled plasma mass spectrometry, and RNA sequencing analyses, *YFH1-off* cells on YPG were inoculated into SC medium at OD_600_ = 1 and grown for 36 hours in a 30°C shaker, diluting to OD_600_ = 1 every 12 hours. After 36 hours, cells were diluted once more to OD_600_ = 1 in either SC or SC-lys-glu and grown for a further 8 hours (44 hours total), or for 5 hours in the experiment in fig. S5B. Cells were then processed for downstream assays.

### Yeast spheroplasts for aconitase assay

Cell density of 44-hr liquid cultures (*YFH1-off* and *H3*^*H113N*^ *YFH1-off*), or exponentially growing cells (WT and *H3*^*H113N*^), was determined and equal amounts of cells from each culture were used for producing spheroplasts (2.5 × 10^9^ cells). Cells were washed once with 50 mL of water and once with 400 µL of DTT Buffer (100 mM Tris-H_2_SO_4_ pH 9.2, 10 mM DTT). The washed cells were resuspended in 400 µL of DTT Buffer and incubated for 20 minutes at 30°C. Treated cells were then washed twice with 1.3 mL of Zymolyase Buffer (20 mM Tris-HCl pH7.5, 1.2 M Sorbitol), resuspended in 1.3 mL Zymolyase Buffer containing 200 µg Zymolyase-100T (Nacalai Tesque), and incubated at 30°C for 35 minutes. Spheroplastization of at least 95% of cells was confirmed under the microscope by a 1:10 dilution into water. Spheroplasts were pelleted at 2000 × g and the supernatant was removed.

### Aconitase assay

Cellular aconitase activity was assessed using the Aconitase Activity Assay Kit (Sigma MAK051-1KT), and the manufacturer’s protocol was followed with minor adjustments. Spheroplasts were lysed in 200 µL of Assay Buffer, centrifuged at 13000 × g for 10 min at 4°C, and the supernatant (i.e., the lysate) was recovered. Protein concentration was determined using the BCA Protein Assay Kit (Pierce) and 900 µg of protein was diluted into a final volume of 100 µL in Assay Buffer containing 1× Protease Inhibitor Cocktail (Pierce). 10 µL of freshly prepared Activator Solution (following the manufacturer’s protocol) was added to the samples and incubated for 1 hour on ice. 50 µL of activated lysate was then mixed with 50 µL of Reaction Mix, which contains either enzyme and substrate, or enzyme alone (for background correction). The reaction was incubated for 1 hour at 25°C, after which 10 µL of developer solution was added to each sample and incubated for another 10 minutes at 25°C. Absorbance at 450 nm was determined, background was subtracted, and the absorbance readings were converted to product concentration (mM of isocitrate) using a standard curve as described by the manufacturer.

### RNA extraction

Cells from the 44-hr cultures (*YFH1-off* and *H3*^*H113N*^ *YFH1-off*), or exponentially growing cells (WT and *H3*^*H113N*^), were collected by centrifugation and frozen at -20°C until processed for RNA extraction and RNA sequencing. RNA was extracted using previously published methods (*71*). RNA extracted for subsequent RNA-seq analysis are from four replicate experiments.

### Sample preparation for poly-A RNA sequencing

Prior to preparing RNA-seq libraries for Illumina NovaSeq sequencing, contaminating DNA was digested using Turbo DNase (Thermo Fisher). RNA quality was then assessed on a 1% agarose gel to confirm minimal RNA degradation. RNA-sequencing libraries were then prepared with the KAPA mRNA Hyper Prep kit (KAPA Biosystems) according to the manufacturer’s protocols. RNA-seq libraries were assessed for correct fragment size and the presence of adapter dimers buy running on a 1% agarose gel. Average library sizes of ∼270 bp were generated. Libraries were pooled for multiplexed sequencing and further purified using KAPA Pure beads. Total DNA concentration was adjusted to 10 nM for Illumina sequencing.

### RNA sequencing and differential gene expression analysis

High throughput sequencing was performed on Illumina’s NovaSeq system. Total read count per library ranged from ∼8-14 million. De-multiplexed reads, in FASTQ file format, were processed to check for read and genome alignment quality, and to convert reads to normalized gene counts (https://training.galaxyproject.org/training-material/topics/transcriptomics/tutorials/rna-seq-reads-to-counts/tutorial.html) (*72*). Processing steps included assessment of raw read quality using FastQC, and genome alignment in strand-specific manner to the R64-1-1 S288C reference genome assembly (sacCer3) using HISAT2 (*73*). Assigning and counting reads for 6692 annotated open reading frames was performed using featureCounts (*74*). Determination of adjusted p-values for differential gene expression comparisons was done using DESeq2 (https://bioconductor.org/packages/release/bioc/vignettes/DESeq2/inst/doc/DESeq2.html) (*75*). Principal component analysis was performed using a built-in function as part of the DESeq2 package. Significantly differentially expressed genes (adjusted p-value <0.05) identified by DESeq2 were assessed for Gene Ontology term enrichment using the Database for Annotation, Visualization, and Integrated Discovery (DAVID) (*76, 77*) with default parameters. Gene sets for gene expression visualization in heatmaps were constructed by downloading gene ontology term gene lists from AmiGO 2 in March 2017.

### Inductively coupled plasma mass spectrometry (ICP-MS)

Cells (8 × 10^8^) from the 44-hr or 41-hr cultures (*YFH1-off* strains), or exponentially growing cells (WT and *H3*^*H113N*^) were collected and washed twice in 5 mM EDTA, to remove cell surface-associated metals, and once in Milli-Q water. Cell pellets were stored at -20°C until further processed for ICP-MS. Cell pellets were overlaid with 143 µL of 70% nitric acid (Fisher, Optima grade, A467-500, Lot 1216040) and digested at room temperature for 24 hours, followed by incubation at 65°C for about 4 hours, and finally being diluted to a final nitric acid concentration of 2% (v/v) with Milli-Q water. Iron and copper contents were determined by ICP-MS on an Agilent 8900 Triple Quadropole ICP-MS instrument, in comparison to an environmental calibration standard (Agilent 5183-4688), using 89Y as an internal standard (Inorganic Ventures MSY-100PPM). The levels of all analytes were determined in MS/MS mode, where 63Cu, was measured directly using He in the collision reaction cell, and 56Fe was directly determined using H_2_ as a cell gas. The average of 3-5 technical replicate measurements was used for each individual biological replicate. The average variation between technical replicate measurements was 1.4% for all analytes and never exceeded 5% for an individual sample. All Cu and Fe measurements were within the calibrated linear ranges and above the lower limits of detection, as determined from multiple blank samples. ICP MassHunter software was used for ICP-MS data analysis.

### Statistical analyses

The number of experimental replicates (n), and the observed significance levels are indicated in figure legends. All statistical analyses were performed using R version 4.0.0, including use of the DESeq2 package, as described above. For the majority of pairwise comparisons, F tests were first performed to determine if group variances were significantly different. When variances were not different, p values for group differences were determined with the two-sample unpaired t-test. When variances were significantly different, p values were determined with the Welch two sample t-test. For pairwise comparisons of ≥2-fold upregulated or downregulated genes (Fig. 3C and D), expression values (normalized read counts) were first averaged across the four biological replicates, transformed into log10 scale, and p values were determined with the paired t-test.

## Acknowledgments

We thank Yong Xue for useful discussions, Leonid Kruglyak and Joshua Bloom for the prototrophic yeast strain, and the UCLA Broad Stem Cell Center Sequencing Core for high-throughput RNA sequencing analysis.

## Author contributions

Conceptualization, O.A.C., N.A., M.V., and S.K.K.; Methodology, O.A.C., N.A., M.V., and S.K.K.; Investigation, O.A.C., N.A., N.V.M., M.V., S.S., and C.C.; Formal Analysis, O.A.C. and M.V.; Writing – Original Draft, O.A.C., N.V.M., and S.K.K.; Writing – Review & Editing, O.A.C., N.A., N.V.M., M.V., S.S., S.S.M. and S.K.K.; Resources, O.A.C., N.A., S.S.M., and S.K.K.; Visualization, O.A.C., and M.V.; Supervision, S.K.K.; Project Administration, S.K.K.; Funding Acquisition, S.S.M., and S.K.K.

## Data and materials availability

All data are available in the main text or the supplementary materials. Gene expression datasets are available on the NCBI GEO database (GSE176575).

## Figures

**Fig. S1.**
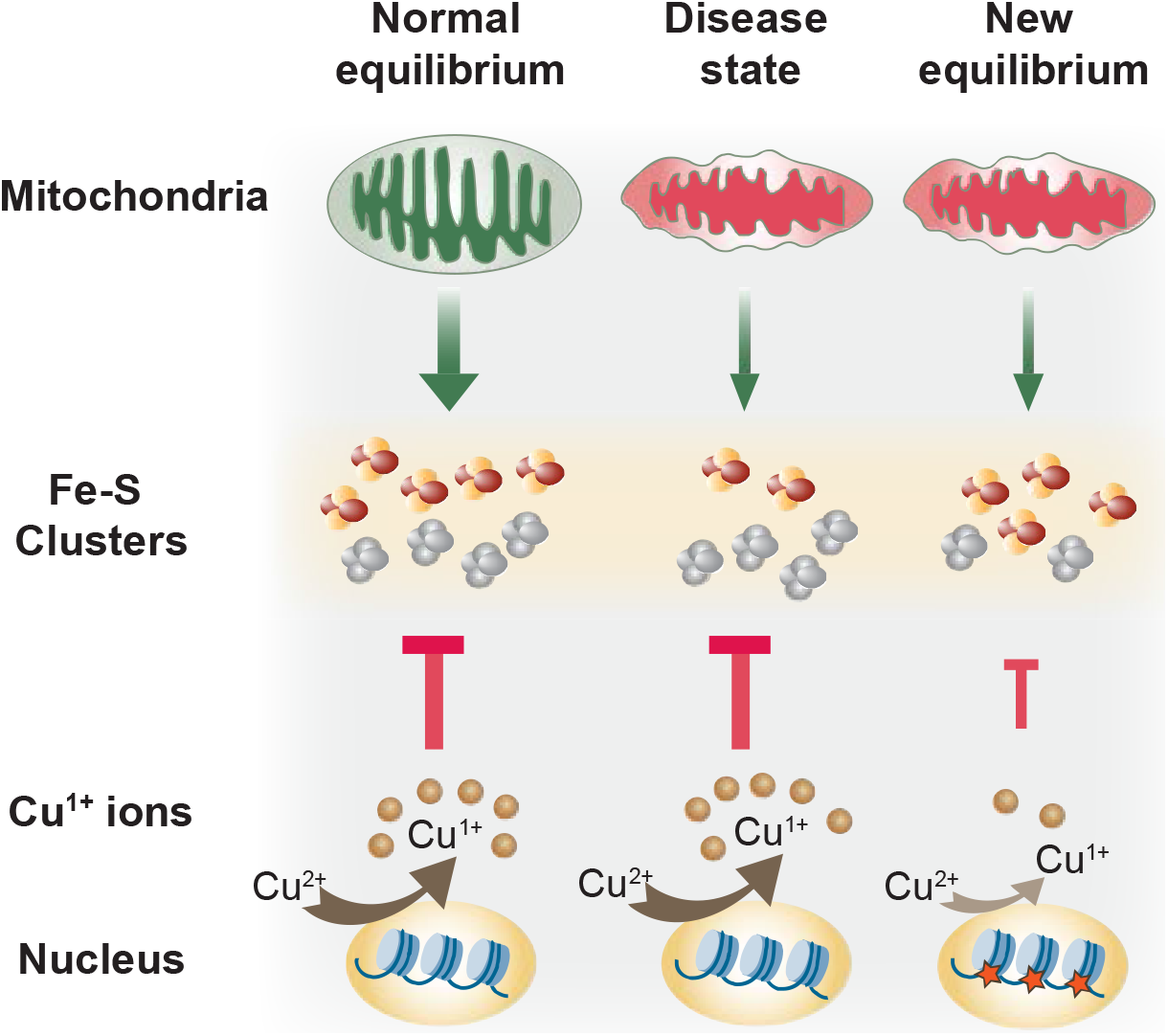
A model depicting the impact of the histone H3 cupric reductase activity and Cu^1+^ on cellular Fe-S cluster quotient. Schematic representation of total Fe-S cluster supply as a balance between Fe-S synthesis and Fe-S attrition due to Cu^1+^ toxicity (left). Inadequate assembly or distribution diminishes the pool of intact Fe-S clusters, making the physiological levels of Cu^1+^ toxic to the cell (middle). Diminishment of Cu^1+^ abundance sets a new equilibrium by decreasing damage to Fe-S clusters, leading to increased quantities despite lower levels of production (right).

**Fig. S2.**
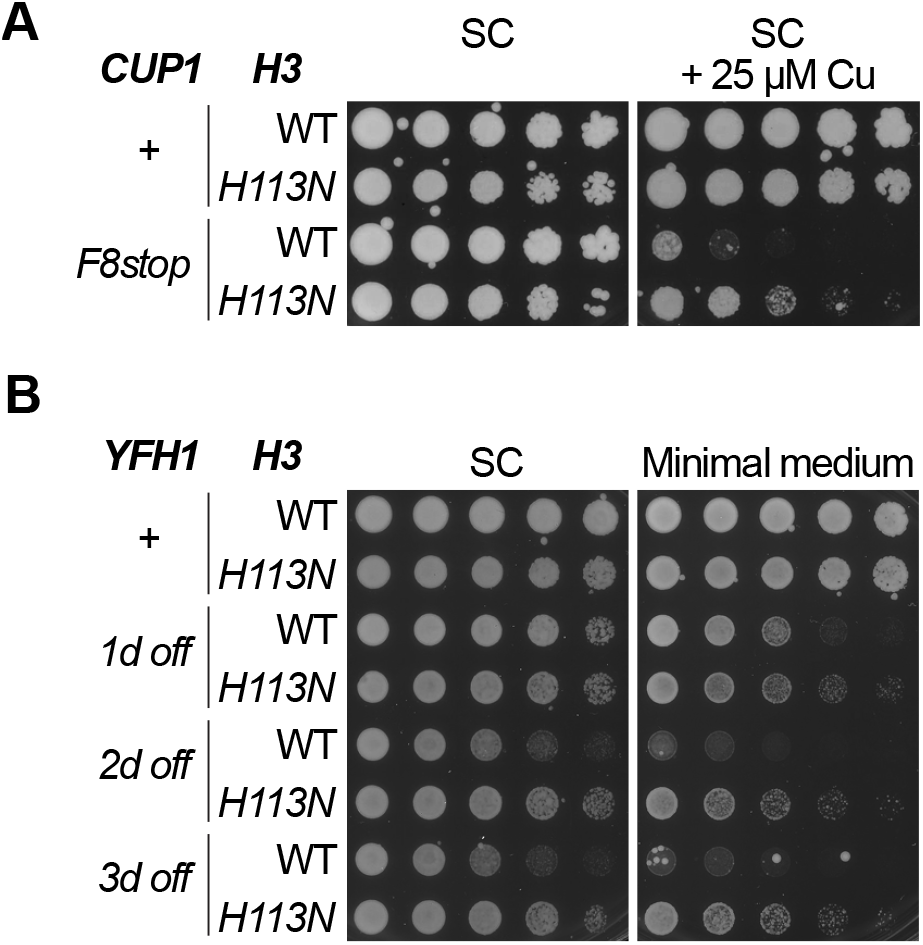
*H3H113N* decreases Cu^1+^ toxicity and rescues the growth defect of *YFH1-off*. (**A and B**) Spot tests with the indicated strains in the BY4741 background (*67*) in fermentative media. Media either contained all twenty amino acids (SC), or lacked all of the amino acids except for leucine, histidine, and methionine (Minimal medium), which the BY4741 strain cannot synthesize. Note that the experiments in Figs. 1C and 2 were performed with a fully prototrophic strain (YLK1879), ruling out a strain-specific phenotype.

**Fig. S3.**
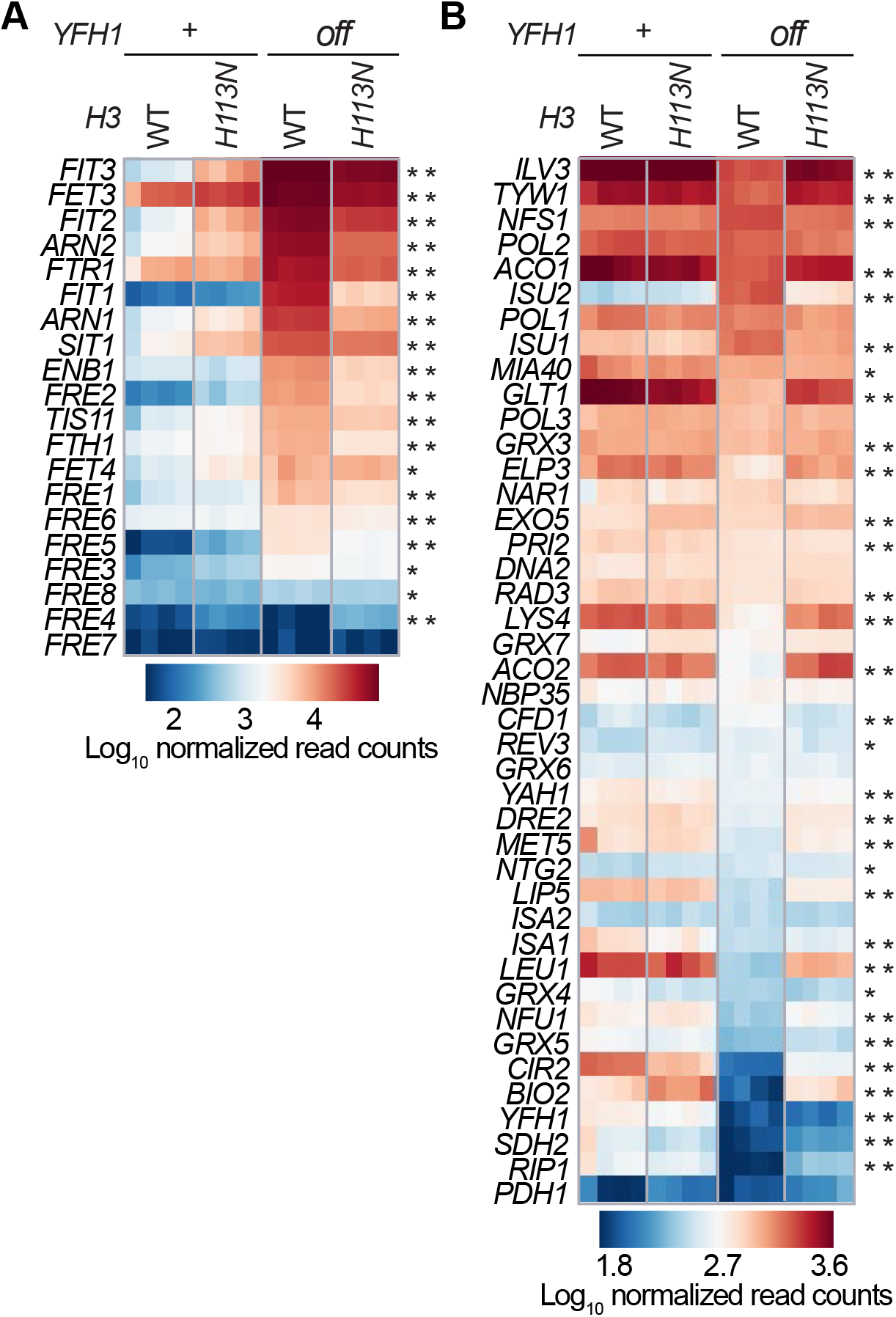
*H3H113N* prevents changes in the iron regulon and Fe-S cluster binding genes. **(A and B)** Gene expression values for (A) iron regulon genes or (B) Fe-S cluster binding genes in the indicated strains growing in SC medium from four independent experiments. *YFH1-off* strains were grown for 44 hrs prior to RNA sequencing. Genes have been ordered based on average expression level in the *YFH1-off* strain. * indicates genes that are significantly differentially expressed (adjusted p < 0.05) in *YFH1-off* compared to WT. ** indicates genes that are significantly differentially expressed (adjusted p < 0.05) in *YFH1-off* compared to WT *<u>and</u> H3*^*H113N*^ *YFH1-off*.

**Fig. S4.**
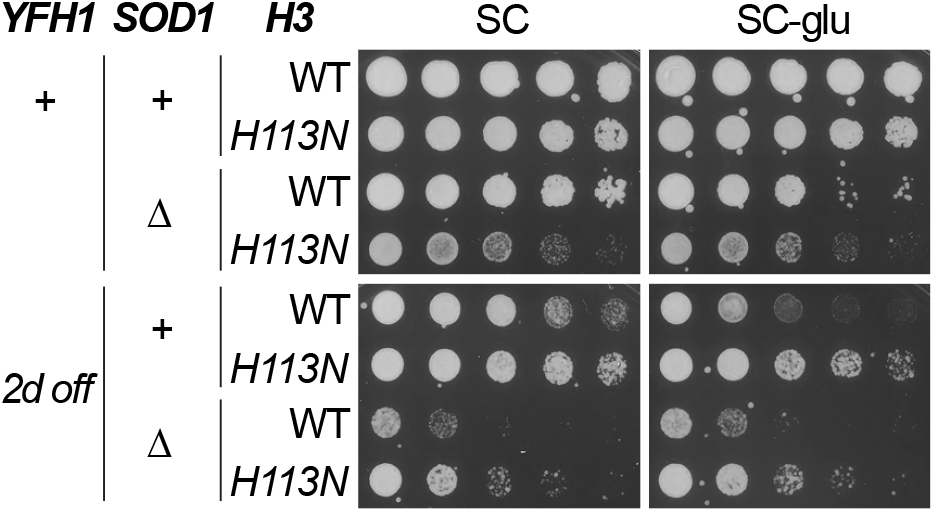
The rescue of YFH1-off by *H3H113N* does not depend on superoxide dismutase activity. Spot test assays in SC medium or SC without glutamate. Note that *sod1Δ* is a lysine auxotroph.

**Fig. S5.**
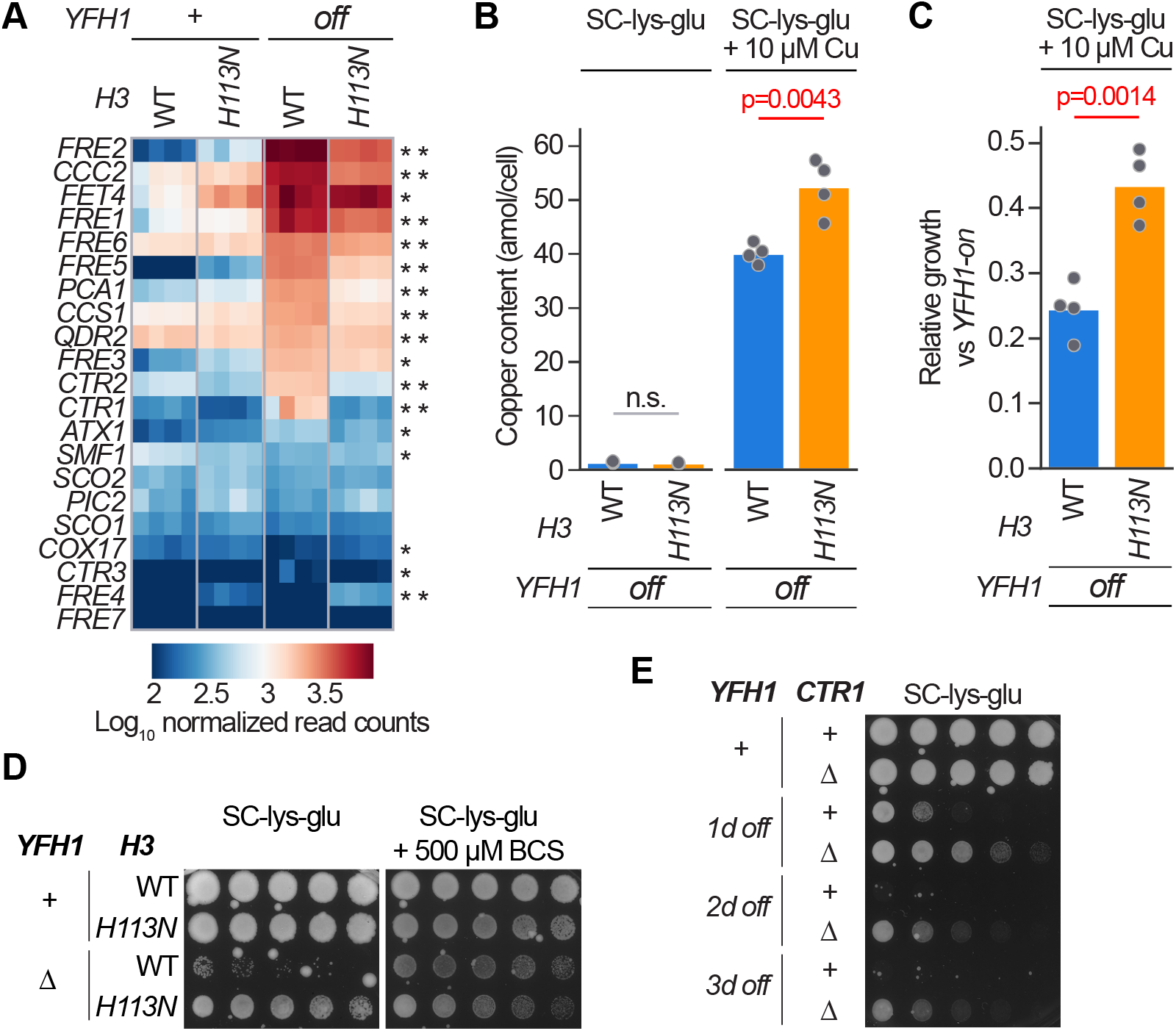
*H3H113N* rescues the loss of Yfh1 function by diminishing Cu^1+^ toxicity. **(A)** Gene expression values for genes involved in copper uptake and mobilization in strains growing in SC medium from four independent experiments. *YFH1-off* strains were grown in SC medium for 44 hours prior to RNA sequencing. Genes have been ordered based on average expression level in the *YFH1-off* strain. * indicates genes that are significantly differentially expressed (adjusted p < 0.05) in *YFH1-off* compared to WT. ** indicates genes that are significantly differentially expressed (adjusted p < 0.05) in *YFH1-off* compared to WT and *H3*^*H113N*^ *YFH1-off*. **(B)** Intracellular copper content for strains grown in liquid SC-lys-glu media ± additional CuSO4. Baseline copper concentration in SC media is ∼0.25 µM. *YFH1-off* strains were grown for 41 hours (see Methods for growth procedure). Bars show means and each dot is an independent experiment (n = 4). n.s. not significant. **(C)** Growth after 41 hours (as in B). Bars show means and each dot is an independent experiment (n = 4). (**D and E**) Spot test assays in SC medium or SC without lysine and glutamate and with or without BCS.

**Table S1.**
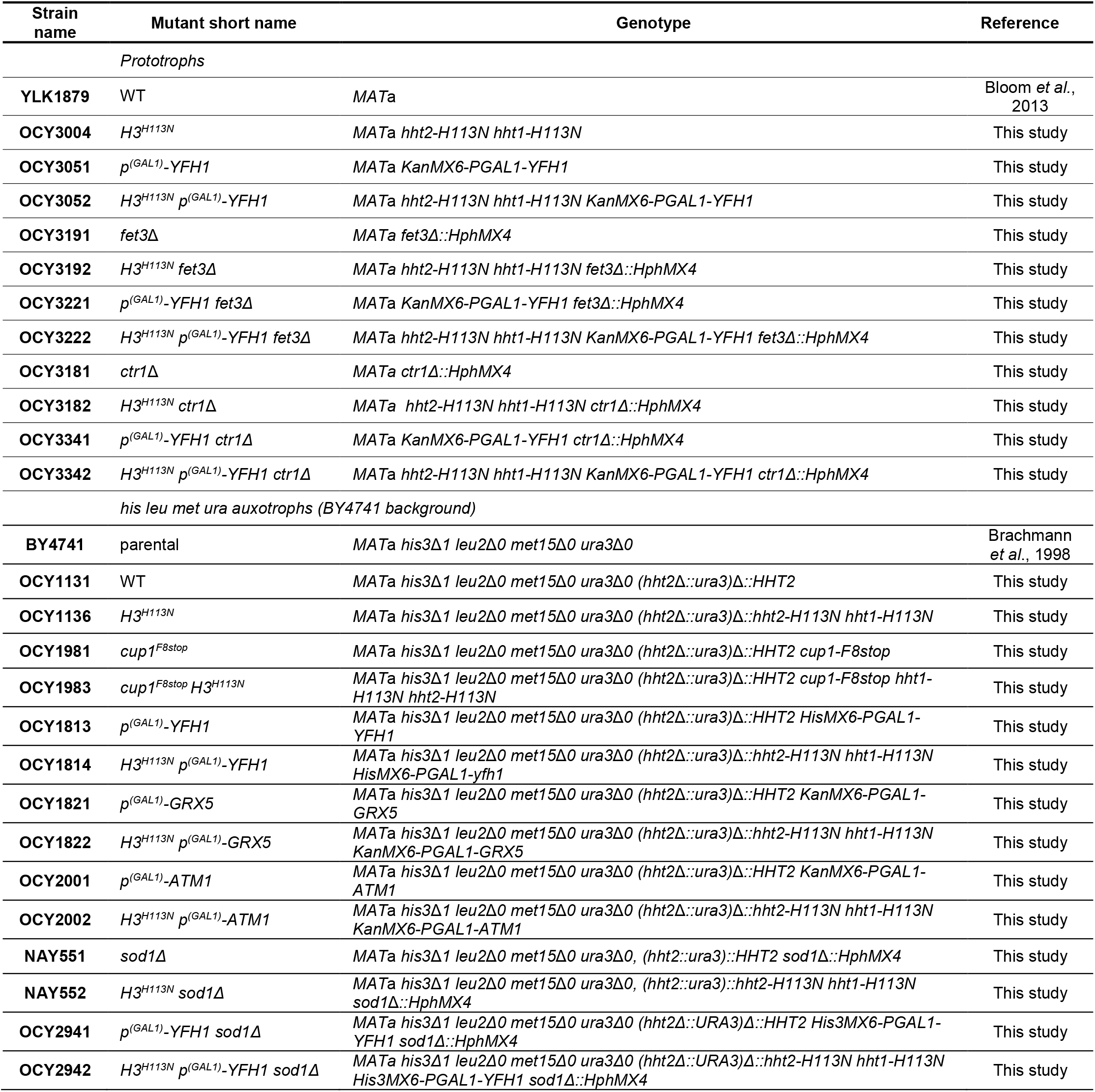
Yeast strains used in this study.

## References

1. H. Beinert, R. H. Holm, E. Munck, Iron-sulfur clusters: nature’s modular, multipurpose structures. Science 277, 653–659 (1997).

2. R. Lill, Function and biogenesis of iron-sulphur proteins. Nature 460, 831–838 (2009).

3. V. Campuzano, L. Montermini, M. D. Molto, L. Pianese, M. Cossee, F. Cavalcanti, E. Monros, F. Rodius, F. Duclos, A. Monticelli, F. Zara, J. Canizares, H. Koutnikova, S. I. Bidichandani, C. Gellera, A. Brice, P. Trouillas, G. De Michele, A. Filla, R. De Frutos, F. Palau, P. I. Patel, S. Di Donato, J. L. Mandel, S. Cocozza, M. Koenig, M. Pandolfo, Friedreich’s ataxia: autosomal recessive disease caused by an intronic GAA triplet repeat expansion. Science 271, 1423–1427 (1996).

4. Y. Zhang, E. R. Lyver, S. A. Knight, D. Pain, E. Lesuisse, A. Dancis, Mrs3p, Mrs4p, and frataxin provide iron for Fe-S cluster synthesis in mitochondria. J Biol Chem 281, 22493–22502 (2006).

5. B. Schilke, C. Voisine, H. Beinert, E. Craig, Evidence for a conserved system for iron metabolism in the mitochondria of Saccharomyces cerevisiae. Proc Natl Acad Sci U S A 96, 10206–10211 (1999).

6. H. Koutnikova, V. Campuzano, F. Foury, P. Dolle, O. Cazzalini, M. Koenig, Studies of human, mouse and yeast homologues indicate a mitochondrial function for frataxin. Nat Genet 16, 345–351 (1997).

7. M. Fontecave, S. Ollagnier-de-Choudens, Iron-sulfur cluster biosynthesis in bacteria: Mechanisms of cluster assembly and transfer. Arch Biochem Biophys 474, 226–237 (2008).

8. U. Muhlenhoff, N. Richhardt, M. Ristow, G. Kispal, R. Lill, The yeast frataxin homolog Yfh1p plays a specific role in the maturation of cellular Fe/S proteins. Hum Mol Genet 11, 2025–2036 (2002).

9. O. S. Chen, S. Hemenway, J. Kaplan, Inhibition of Fe-S cluster biosynthesis decreases mitochondrial iron export: evidence that Yfh1p affects Fe-S cluster synthesis. Proc Natl Acad Sci U S A 99, 12321–12326 (2002).

10. A. Rotig, P. de Lonlay, D. Chretien, F. Foury, M. Koenig, D. Sidi, A. Munnich, P. Rustin, Aconitase and mitochondrial iron-sulphur protein deficiency in Friedreich ataxia. Nat Genet 17, 215–217 (1997).

11. A. L. Bulteau, H. A. O’Neill, M. C. Kennedy, M. Ikeda-Saito, G. Isaya, L. I. Szweda, Frataxin acts as an iron chaperone protein to modulate mitochondrial aconitase activity. Science 305, 242–245 (2004).

12. F. Fazius, E. Shelest, P. Gebhardt, M. Brock, The fungal alpha-aminoadipate pathway for lysine biosynthesis requires two enzymes of the aconitase family for the isomerization of homocitrate to homoisocitrate. Mol Microbiol 86, 1508–1530 (2012).

13. X. J. Chen, X. Wang, B. A. Kaufman, R. A. Butow, Aconitase couples metabolic regulation to mitochondrial DNA maintenance. Science 307, 714–717 (2005).

14. S. Jang, J. A. Imlay, Micromolar intracellular hydrogen peroxide disrupts metabolism by damaging iron-sulfur enzymes. J Biol Chem 282, 929–937 (2007).

15. E. S. Boyd, K. M. Thomas, Y. Dai, J. M. Boyd, F. W. Outten, Interplay between oxygen and Fe-S cluster biogenesis: insights from the Suf pathway. Biochemistry 53, 5834–5847 (2014).

16. J. A. Imlay, Cellular defenses against superoxide and hydrogen peroxide. Annu Rev Biochem 77, 755–776 (2008).

17. G. Tan, J. Yang, T. Li, J. Zhao, S. Sun, X. Li, C. Lin, J. Li, H. Zhou, J. Lyu, H. Ding, Anaerobic Copper Toxicity and Iron-Sulfur Cluster Biogenesis in Escherichia coli. Appl Environ Microbiol 83, (2017).

18. A. Shanmuganathan, S. V. Avery, S. A. Willetts, J. E. Houghton, Copper-induced oxidative stress in Saccharomyces cerevisiae targets enzymes of the glycolytic pathway. FEBS Lett 556, 253–259 (2004).

19. S. V. Avery, N. G. Howlett, S. Radice, Copper toxicity towards Saccharomyces cerevisiae: dependence on plasma membrane fatty acid composition. Appl Environ Microbiol 62, 3960–3966 (1996).

20. N. J. Robinson, D. R. Winge, Copper metallochaperones. Annu Rev Biochem 79, 537–562 (2010).

21. D. Brancaccio, A. Gallo, M. Piccioli, E. Novellino, S. Ciofi-Baffoni, L. Banci, [4Fe-4S] Cluster Assembly in Mitochondria and Its Impairment by Copper. J Am Chem Soc 139, 719–730 (2017).

22. L. Macomber, J. A. Imlay, The iron-sulfur clusters of dehydratases are primary intracellular targets of copper toxicity. Proc Natl Acad Sci U S A 106, 8344–8349 (2009).

23. C. Vallieres, S. L. Holland, S. V. Avery, Mitochondrial Ferredoxin Determines Vulnerability of Cells to Copper Excess. Cell Chem Biol 24, 1228–1237 e1223 (2017).

24. N. Attar, O. A. Campos, M. Vogelauer, C. Cheng, Y. Xue, S. Schmollinger, L. Salwinski, N. V. Mallipeddi, B. A. Boone, L. Yen, S. Yang, S. Zikovich, J. Dardine, M. F. Carey, S. S. Merchant, S. K. Kurdistani, The histone H3-H4 tetramer is a copper reductase enzyme. Science 369, 59–64 (2020).

25. D. R. Winge, K. B. Nielson, W. R. Gray, D. H. Hamer, Yeast metallothionein. Sequence and metal-binding properties. J Biol Chem 260, 14464–14470 (1985).

26. S. Fogel, J. W. Welch, Tandem gene amplification mediates copper resistance in yeast. Proc Natl Acad Sci U S A 79, 5342–5346 (1982).

27. G. Karthikeyan, L. K. Lewis, M. A. Resnick, The mitochondrial protein frataxin prevents nuclear damage. Hum Mol Genet 11, 1351–1362 (2002).

28. M. T. Rodriguez-Manzaneque, J. Tamarit, G. Belli, J. Ros, E. Herrero, Grx5 is a mitochondrial glutaredoxin required for the activity of iron/sulfur enzymes. Mol Biol Cell 13, 1109–1121 (2002).

29. M. A. Uzarska, R. Dutkiewicz, S. A. Freibert, R. Lill, U. Muhlenhoff, The mitochondrial Hsp70 chaperone Ssq1 facilitates Fe/S cluster transfer from Isu1 to Grx5 by complex formation. Mol Biol Cell 24, 1830–1841 (2013).

30. G. Kispal, P. Csere, C. Prohl, R. Lill, The mitochondrial proteins Atm1p and Nfs1p are essential for biogenesis of cytosolic Fe/S proteins. EMBO J 18, 3981–3989 (1999).

31. R. E. Miller, E. R. Stadtman, Glutamate synthase from Escherichia coli. An iron-sulfide flavoprotein. J Biol Chem 247, 7407–7419 (1972).

32. U. Muhlenhoff, N. Richter, O. Pines, A. J. Pierik, R. Lill, Specialized function of yeast Isa1 and Isa2 proteins in the maturation of mitochondrial [4Fe-4S] proteins. J Biol Chem 286, 41205–41216 (2011).

33. B. R. Crane, L. M. Siegel, E. D. Getzoff, Sulfite reductase structure at 1.6 A: evolution and catalysis for reduction of inorganic anions. Science 270, 59–67 (1995).

34. S. C. Kennedy, R. Rauner, O. Gawron, On pig heart aconitase. Biochem Biophys Res Commun 47, 740–745 (1972).

35. F. J. Ruzicka, H. Beinert, The soluble “high potential” type iron-sulfur protein from mitochondria is aconitase. J Biol Chem 253, 2514–2517 (1978).

36. A. H. Robbins, C. D. Stout, Structure of activated aconitase: formation of the [4Fe-4S] cluster in the crystal. Proc Natl Acad Sci U S A 86, 3639–3643 (1989).

37. F. Foury, D. Talibi, Mitochondrial control of iron homeostasis. A genome wide analysis of gene expression in a yeast frataxin-deficient strain. J Biol Chem 276, 7762–7768 (2001).

38. Y. Yamaguchi-Iwai, A. Dancis, R. D. Klausner, AFT1: a mediator of iron regulated transcriptional control in Saccharomyces cerevisiae. EMBO J 14, 1231–1239 (1995).

39. O. S. Chen, R. J. Crisp, M. Valachovic, M. Bard, D. R. Winge, J. Kaplan, Transcription of the yeast iron regulon does not respond directly to iron but rather to iron-sulfur cluster biosynthesis. J Biol Chem 279, 29513–29518 (2004).

40. A. Kumanovics, O. S. Chen, L. Li, D. Bagley, E. M. Adkins, H. Lin, N. N. Dingra, C. E. Outten, G. Keller, D. Winge, D. M. Ward, J. Kaplan, Identification of FRA1 and FRA2 as genes involved in regulating the yeast iron regulon in response to decreased mitochondrial iron-sulfur cluster synthesis. J Biol Chem 283, 10276–10286 (2008).

41. S. Puig, E. Askeland, D. J. Thiele, Coordinated remodeling of cellular metabolism during iron deficiency through targeted mRNA degradation. Cell 120, 99–110 (2005).

42. S. Puig, S. V. Vergara, D. J. Thiele, Cooperation of two mRNA-binding proteins drives metabolic adaptation to iron deficiency. Cell Metab 7, 555–564 (2008).

43. O. Protchenko, T. Ferea, J. Rashford, J. Tiedeman, P. O. Brown, D. Botstein, C. C. Philpott, Three cell wall mannoproteins facilitate the uptake of iron in Saccharomyces cerevisiae. J Biol Chem 276, 49244–49250 (2001).

44. C. W. Yun, T. Ferea, J. Rashford, O. Ardon, P. O. Brown, D. Botstein, J. Kaplan, C. C. Philpott, Desferrioxamine-mediated iron uptake in Saccharomyces cerevisiae. Evidence for two pathways of iron uptake. J Biol Chem 275, 10709–10715 (2000).

45. F. Foury, O. Cazzalini, Deletion of the yeast homologue of the human gene associated with Friedreich’s ataxia elicits iron accumulation in mitochondria. FEBS Lett 411, 373–377 (1997).

46. M. Babcock, D. de Silva, R. Oaks, S. Davis-Kaplan, S. Jiralerspong, L. Montermini, M. Pandolfo, J. Kaplan, Regulation of mitochondrial iron accumulation by Yfh1p, a putative homolog of frataxin. Science 276, 1709–1712 (1997).

47. M. B. Delatycki, J. Camakaris, H. Brooks, T. Evans-Whipp, D. R. Thorburn, R. Williamson, S. M. Forrest, Direct evidence that mitochondrial iron accumulation occurs in Friedreich ataxia. Ann Neurol 45, 673–675 (1999).

48. R. Stearman, D. S. Yuan, Y. Yamaguchi-Iwai, R. D. Klausner, A. Dancis, A permease-oxidase complex involved in high-affinity iron uptake in yeast. Science 271, 1552–1557 (1996).

49. C. Askwith, D. Eide, A. Van Ho, P. S. Bernard, L. Li, S. Davis-Kaplan, D. M. Sipe, J. Kaplan, The FET3 gene of S. cerevisiae encodes a multicopper oxidase required for ferrous iron uptake. Cell 76, 403–410 (1994).

50. Y. Yamaguchi-Iwai, R. Stearman, A. Dancis, R. D. Klausner, Iron-regulated DNA binding by the AFT1 protein controls the iron regulon in yeast. EMBO J 15, 3377–3384 (1996).

51. J. M. De Freitas, A. Liba, R. Meneghini, J. S. Valentine, E. B. Gralla, Yeast lacking Cu-Zn superoxide dismutase show altered iron homeostasis. Role of oxidative stress in iron metabolism. J Biol Chem 275, 11645–11649 (2000).

52. S. J. Lin, V. C. Culotta, Suppression of oxidative damage by Saccharomyces cerevisiae ATX2, which encodes a manganese-trafficking protein that localizes to Golgi-like vesicles. Mol Cell Biol 16, 6303–6312 (1996).

53. A. Dancis, D. Haile, D. S. Yuan, R. D. Klausner, The Saccharomyces cerevisiae copper transport protein (Ctr1p). Biochemical characterization, regulation by copper, and physiologic role in copper uptake. J Biol Chem 269, 25660–25667 (1994).

54. K. D. MacIsaac, T. Wang, D. B. Gordon, D. K. Gifford, G. D. Stormo, E. Fraenkel, An improved map of conserved regulatory sites for Saccharomyces cerevisiae. BMC Bioinformatics 7, 113 (2006).

55. M. Darash-Yahana, Y. Pozniak, M. Y. Lu, Y. S. Sohn, O. Karmi, S. Tamir, F. Bai, L. H. Song, P. A. Jennings, E. Pikarsky, T. Geiger, J. N. Onuchic, R. Mittler, R. Nechushtai, Breast cancer tumorigenicity is dependent on high expression levels of NAF-1 and the lability of its Fe-S clusters. P Natl Acad Sci USA 113, 10890–10895 (2016).

56. S. Oshiro, M. S. Morioka, M. Kikuchi, Dysregulation of iron metabolism in Alzheimer’s disease, Parkinson’s disease, and amyotrophic lateral sclerosis. Adv Pharmacol Sci 2011, 378278 (2011).

57. T. A. Rouault, W. H. Tong, Iron-sulfur cluster biogenesis and human disease. Trends Genet 24, 398–407 (2008).

58. J. Rudolf, V. Makrantoni, W. J. Ingledew, M. J. R. Stark, M. F. White, The DNA repair helicases XPD and FancJ have essential iron-sulfur domains. Mol Cell 23, 801–808 (2006).

59. R. Jain, E. S. Vanamee, B. G. Dzikovski, A. Buku, R. E. Johnson, L. Prakash, S. Prakash, A. K. Aggarwal, An iron-sulfur cluster in the polymerase domain of yeast DNA polymerase epsilon. J Mol Biol 426, 301–308 (2014).

60. H. Ye, S. Y. Jeong, M. C. Ghosh, G. Kovtunovych, L. Silvestri, D. Ortillo, N. Uchida, J. Tisdale, C. Camaschella, T. A. Rouault, Glutaredoxin 5 deficiency causes sideroblastic anemia by specifically impairing heme biosynthesis and depleting cytosolic iron in human erythroblasts. J Clin Invest 120, 1749–1761 (2010).

61. J. Boultwood, A. Pellagatti, M. Nikpour, B. Pushkaran, C. Fidler, H. Cattan, T. J. Littlewood, L. Malcovati, M. G. Della Porta, M. Jadersten, S. Killick, A. Giagounidis, D. Bowen, E. Hellstrom-Lindberg, M. Cazzola, J. S. Wainscoat, The role of the iron transporter ABCB7 in refractory anemia with ring sideroblasts. PLoS One 3, e1970 (2008).

62. R. A. Vaubel, G. Isaya, Iron-sulfur cluster synthesis, iron homeostasis and oxidative stress in Friedreich ataxia. Mol Cell Neurosci 55, 50–61 (2013).

63. T. Ast, J. D. Meisel, S. Patra, H. Wang, R. M. H. Grange, S. H. Kim, S. E. Calvo, L. L. Orefice, F. Nagashima, F. Ichinose, W. M. Zapol, G. Ruvkun, D. P. Barondeau, V. K. Mootha, Hypoxia Rescues Frataxin Loss by Restoring Iron Sulfur Cluster Biogenesis. Cell 177, 1507–1521 e1516 (2019).

64. A. K. Pandey, J. Pain, A. Dancis, D. Pain, Mitochondria export iron-sulfur and sulfur intermediates to the cytoplasm for iron-sulfur cluster assembly and tRNA thiolation in yeast. J Biol Chem 294, 9489–9502 (2019).

65. S. A. Pearson, C. Wachnowsky, J. A. Cowan, Defining the mechanism of the mitochondrial Atm1p [2Fe-2S] cluster exporter. Metallomics 12, 902–915 (2020).

66. R. Allikmets, W. H. Raskind, A. Hutchinson, N. D. Schueck, M. Dean, D. M. Koeller, Mutation of a putative mitochondrial iron transporter gene (ABC7) in X-linked sideroblastic anemia and ataxia (XLSA/A). Hum Mol Genet 8, 743–749 (1999).

67. C. B. Brachmann, A. Davies, G. J. Cost, E. Caputo, J. Li, P. Hieter, J. D. Boeke, Designer deletion strains derived from Saccharomyces cerevisiae S288C: a useful set of strains and plasmids for PCR-mediated gene disruption and other applications. Yeast 14, 115–132 (1998).

68. J. S. Bloom, I. M. Ehrenreich, W. T. Loo, T. L. Lite, L. Kruglyak, Finding the sources of missing heritability in a yeast cross. Nature 494, 234–237 (2013).

69. O. W. Ryan, J. M. Skerker, M. J. Maurer, X. Li, J. C. Tsai, S. Poddar, M. E. Lee, W. DeLoache, J. E. Dueber, A. P. Arkin, J. H. Cate, Selection of chromosomal DNA libraries using a multiplex CRISPR system. Elife 3, (2014).

70. M. S. Longtine, A. McKenzie, 3rd, D. J. Demarini, N. G. Shah, A. Wach, A. Brachat, P. Philippsen, J. R. Pringle, Additional modules for versatile and economical PCR-based gene deletion and modification in Saccharomyces cerevisiae. Yeast 14, 953–961 (1998).

71. M. E. Schmitt, T. A. Brown, B. L. Trumpower, A rapid and simple method for preparation of RNA from Saccharomyces cerevisiae. Nucleic Acids Res 18, 3091–3092 (1990).

72. B. Batut, S. Hiltemann, A. Bagnacani, D. Baker, V. Bhardwaj, C. Blank, A. Bretaudeau, L. Brillet-Gueguen, M. Cech, J. Chilton, D. Clements, O. Doppelt-Azeroual, A. Erxleben, M. A. Freeberg, S. Gladman, Y. Hoogstrate, H. R. Hotz, T. Houwaart, P. Jagtap, D. Lariviere, G. Le Corguille, T. Manke, F. Mareuil, F. Ramirez, D. Ryan, F. C. Sigloch, N. Soranzo, J. Wolff, P. Videm, M. Wolfien, A. Wubuli, D. Yusuf, N. Galaxy Training, J. Taylor, R. Backofen, A. Nekrutenko, B. Gruning, Community-Driven Data Analysis Training for Biology. Cell Syst 6, 752–758 e751 (2018).

73. D. Kim, J. M. Paggi, C. Park, C. Bennett, S. L. Salzberg, Graph-based genome alignment and genotyping with HISAT2 and HISAT-genotype. Nat Biotechnol 37, 907–915 (2019).

74. Y. Liao, G. K. Smyth, W. Shi, featureCounts: an efficient general purpose program for assigning sequence reads to genomic features. Bioinformatics 30, 923–930 (2014).

75. M. I. Love, W. Huber, S. Anders, Moderated estimation of fold change and dispersion for RNA-seq data with DESeq2. Genome Biol 15, 550 (2014).

76. D. W. Huang, B. T. Sherman, R. A. Lempicki, Systematic and integrative analysis of large gene lists using DAVID bioinformatics resources. Nat Protoc 4, 44–57 (2009).

77. D. W. Huang, B. T. Sherman, R. A. Lempicki, Bioinformatics enrichment tools: paths toward the comprehensive functional analysis of large gene lists. Nucleic Acids Res 37, 1–13 (2009).

